# Natural genetic and transcriptomic variation in photosynthesis associated pathways contribute to adaptive traits variation in worldwide *Arabidopsis thaliana* population

**DOI:** 10.1101/2024.08.29.610265

**Authors:** Wei Liu, Jing Hou, Li Liu, Mengyu Lei, Yu Han, Mingjia Zhu, Wenjia Zhang, Ran Hao, Yan Ji, Huan Si, Jianquan Liu, Yanjun Zan

**Affiliations:** Key Laboratory for Bio-Resource and Eco-Environment of Ministry of Education & Sichuan Zoige Alpine Wetland Ecosystem National Observation and Research Station, College of Life Science, Sichuan University, Chengdu, CN-610065, China; Tobacco Research Institute, Chinese Academy of Agricultural Sciences, Qingdao, CN- 266000, China; State Key Laboratory of Grassland Agro-ecosystem, College of Ecology, Lanzhou University, Lanzhou, CN-730000, China

**Keywords:** *Arabidopsis thaliana*, photosynthesis pathways, natural variation, adaptive traits, crop breeding

## Abstract

Photosynthesis is the most important reaction underlying carbon fixation. Despite its potential in boosting carbon assimilation, nature variations underlying genes in photosynthesis pathway and their role in adaptive traits variation remains elusive. In this study, we investigated the genetic, transcriptomic variation of 1103 genes in photosynthesis associated pathways, including 82 photosynthesis core genes, 24 plastid-encoded RNA polymerase related genes, 2 nucleus-encoded RNA polymerase-related genes, 34 photomorphogenesis-related genes, 40 genes involved in transcription and translation (TAC) and 938 other nuclear-encoded chloroplast-targeted genes. By *de novo* assembling the chloroplast genomes of 28 representative accessions and leveraging whole-genome, transcriptome sequencing data from the 1001 Genome Project, we revealed extensive natural genetic and transcriptome variations these genes in worldwide *Arabidopsis thaliana* population. 34.0% of them were identified with regulatory variations in expression quantitative locus mapping (eQTL) mapping, including key components of Rubisco (*RBCS1B, RBCS2B*), and Rubisco activase (*RCA*). Genome-wide and transcriptome-wide association analysis (GWAS/TWAS) showed that these genetic and transcriptomic variations made considerable contribution to variation of adaptive traits. Overall, our study provides insight into the natural genetic variation of these genes among worldwide *Arabidopsis thaliana* accessions and their role in complex traits variation and adaptation.

## Introduction

Photosynthesis is a complex biochemical process that assimilates radiation to produce sugars for plant growth and fruit setting^1-4^. With the growing demand for food production, optimizing photosynthesis may present new opportunities for crop improvement and breeding. Previous studies on boosting photosynthesis activity have focused on two approaches. The first approach was modifying well-characterized photosynthetic genes. For example, modifying Ribulose-1,5-bisphosphate carboxylase/oxygenase (RuBisCO) and other photosynthesis-related genes^5-7^ to increase carbon assimilation efficacy^8,9^. Developing thermotolerant RuBisCO activase (RCA) to sustain RuBisCO activation under extreme temperature^10,11^ and optimizing light reactions and energy portioning in light harvesting^12,13^. Reducing the variation of photosynthetic induction to increase photosynthetic efficiency in fluctuating light^14-16^. The second approach focused on identifying natural genetic variants with potential for boosting photosynthesis activity ^17,18^. For instance, previous studies have identified quantitative trait loci (QTLs) for photosynthetic activity in various plants, including *Arabidopsis thaliana*, wheat, rice, maize, and tomato^19-27^. Although initial attempt to introduce these QTLs into rice cultivars only enhance overall biomass but not yield ^18^, these discoveries suggest that there is natural genetic variation within species that impacts photosynthetic activity.

While reports showed that artificial selection has improved photosynthesis activity, enhancing photosynthesis has not yet been a primary goal for crop breeding programs^28-30^. This is because the genetic basis of photosynthesis remains elusive^31^. There is a lack of understanding of natural variation in genes involved in photosynthesis, their regulatory polymorphisms, and assessment of their role in complex traits variation. Taking these facts together, plant photosynthesis presents an underutilized reservoir for improving carbon assimilation. These factors highlight the necessity for a holistic investigation of the genetic diversity in photosynthesis associated pathways and their potential in future crop and tree breeding.

In this study, we investigated the genetic and transcriptomic variation of 1103 genes involved in photosynthesis pathway, photomorphogenesis, transcription and translation process of chloroplast, including photosynthesis-core genes (Core), plastid-encoded RNA polymerase related genes (PEP), nucleus-encoded RNA polymerase-related genes (NEP), photomorphogenesis-related genes (PMPG), genes involved in the process of transcription and translation(TAC) and other nuclear-encoded chloroplast-targeted genes (ONECTG,Table S4 and S5). We revealed extensive natural genetic and transcriptome variation for these genes by assembling the chloroplast genomes of 28 representative accessions and leveraging whole genomes, transcriptomes sequencing data from the 1001 Genome Project. By performing expression quantitative locus mapping (eQTL), genome-wide and transcriptome-wide association analysis (GWAS/TWAS), we revealed important contribution from natural genetic and transcriptome variation of photosynthesis genes to variation of adaptive traits and found a number of regulatory variations for key photosynthesis genes. For example, we detected three eQTL affected the expression of *RBCS1B, RBCS2B*, and *RCA*. One eQTL associated with *RCA* expression was significantly associated with fruit number and elevation, highlighting the contribution to adaptive traits variation. Overall, our study provides insight into the natural genetic variation of these genes among worldwide *Arabidopsis thaliana* accessions and their role in complex traits variation and adaptation.

## Results

### 28 *de novo* assembled chloroplast genomes reveal abundant genetic diversity in the *Arabidopsis thaliana* chloroplast genome

To identify natural genetic variation in genes from photosynthesis associated pathways, we *de novo* assembled the chloroplast genomes of 28 worldwide *Arabidopsis thaliana* accessions (Fig 1A and 1C) using Pacbio Sequel II reads. Taking the Col-0 ecotype together, these 29 accessions included both relict and non-relict lines covering a wide range of ecological habitats. The length of assembled chloroplast genomes varied from 154275 to 154528 bp (Table S1 and S2, Fig 1B) with 4.4% regions including *psaJ, rbcL* and *rps16* showed elevated nucleotide diversity (*Pi* value > 0.002, Fig 1D, Fig S1, Table S3). Those results demonstrated extensive genetic diversity in the *Arabidopsis thaliana* chloroplast genome.

**Fig 1.**
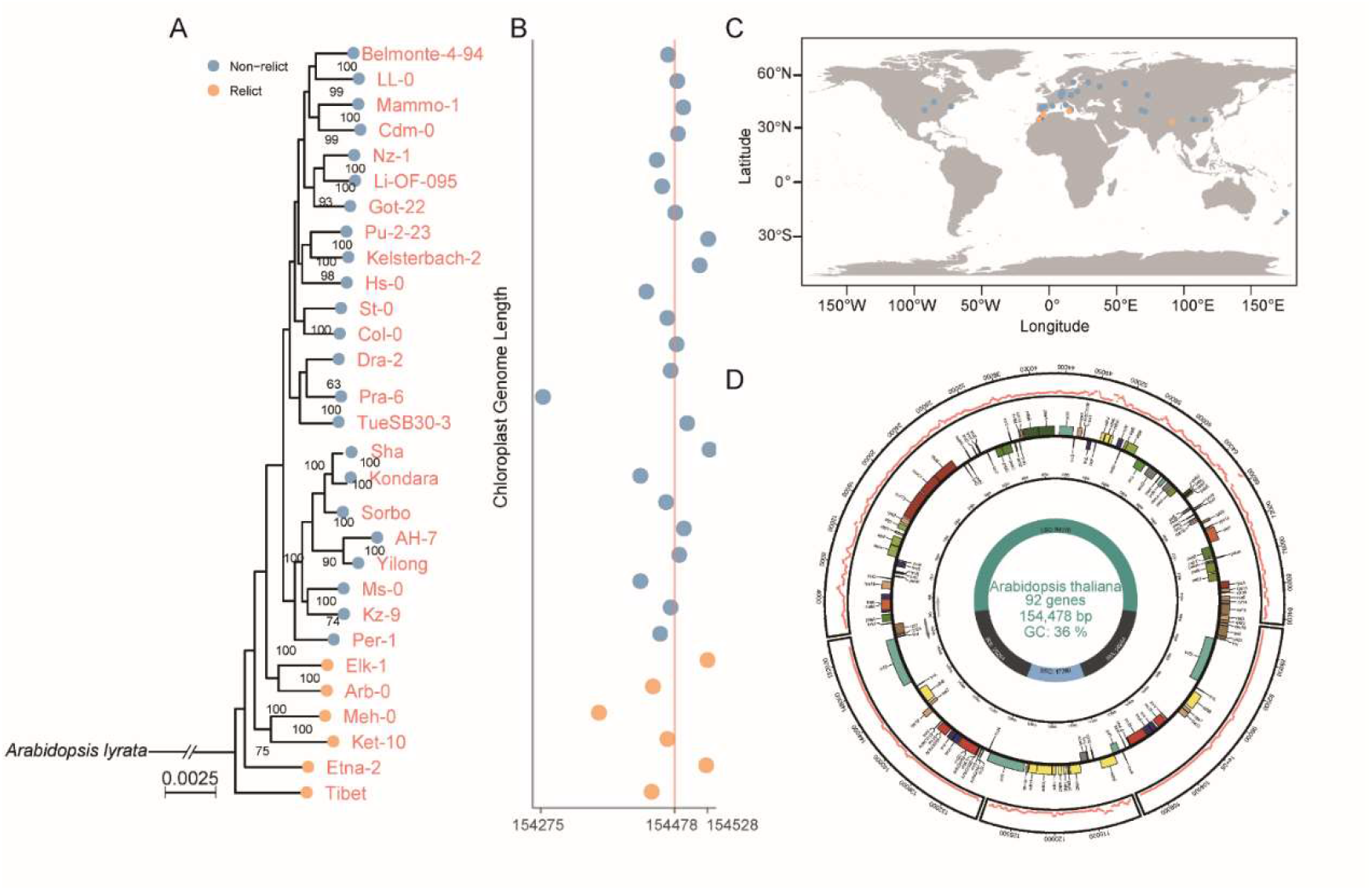
28 *de novo* assembled chloroplast genomes reveal abundant genetic diversity in the *Arabidopsis thaliana* chloroplast genome. (A) A phylogenetic tree of 29 *Arabidopsis thaliana accessions*. (B) Length of the assembled chloroplast genomes. (C) Geographical distribution of these accessions. (D) Distribution of nucleotide diversity (*Pi*, outer layer) along 29 chloroplast genomes.

### Rich genetic diversity in nuclear and plastid-encoded genes (ANPEG)

To investigate genetic diversity of genes involved in photosynthesis associated pathways, we extracted single nucleotide polymorphisms (SNP), insertions/deletions, and structure variations (SVs) for ANPEG, including Core, PEP, NEP, PMPG, TAC and ONECTG (Fig 2A, Table S4 and S5). Overall, these genes were widely distributed across the five chromosomes (Fig. S2). Compared to the average genetic diversity of the genome (background), there are no significant results in nuclear and plastid-encoded genes indicating the genetic diversity of these genes is at the average level (Fig 2B and 2C, Table S6 and S7). In addition, we discovered structural variations (SVs) and Insertions/Deletions (InDels) in gene body of several photosynthesis-core genes: *AT4G02770 (PSAD-1), AT1G32550* (FdC2), *atpF, petB, psbA, rbc*, etc. several PEP components : *AT1G64860 (SIGA), AT1G08540* (*SIGB*) etc., several TAC genes : *AT3G20540 (POLGAMMA1), AT5G04590 (SIR)* etc., several PMPG genes : *AT2G20180 (PIL5), AT1G09530 (PIF3)* etc. A number of genes, such as *rbcL, psbA, psbJ, AT4G04640 (ATPC1)*, and *AT3G50820 (PSBO-2)* etc., were surrounded by high-frequency SVs or InDels (Fig 2D and 2E, Table S8, S9 and S10). Overall, these results demonstrated abundant natural genetic variation for ANPEG.

**Fig 2.**
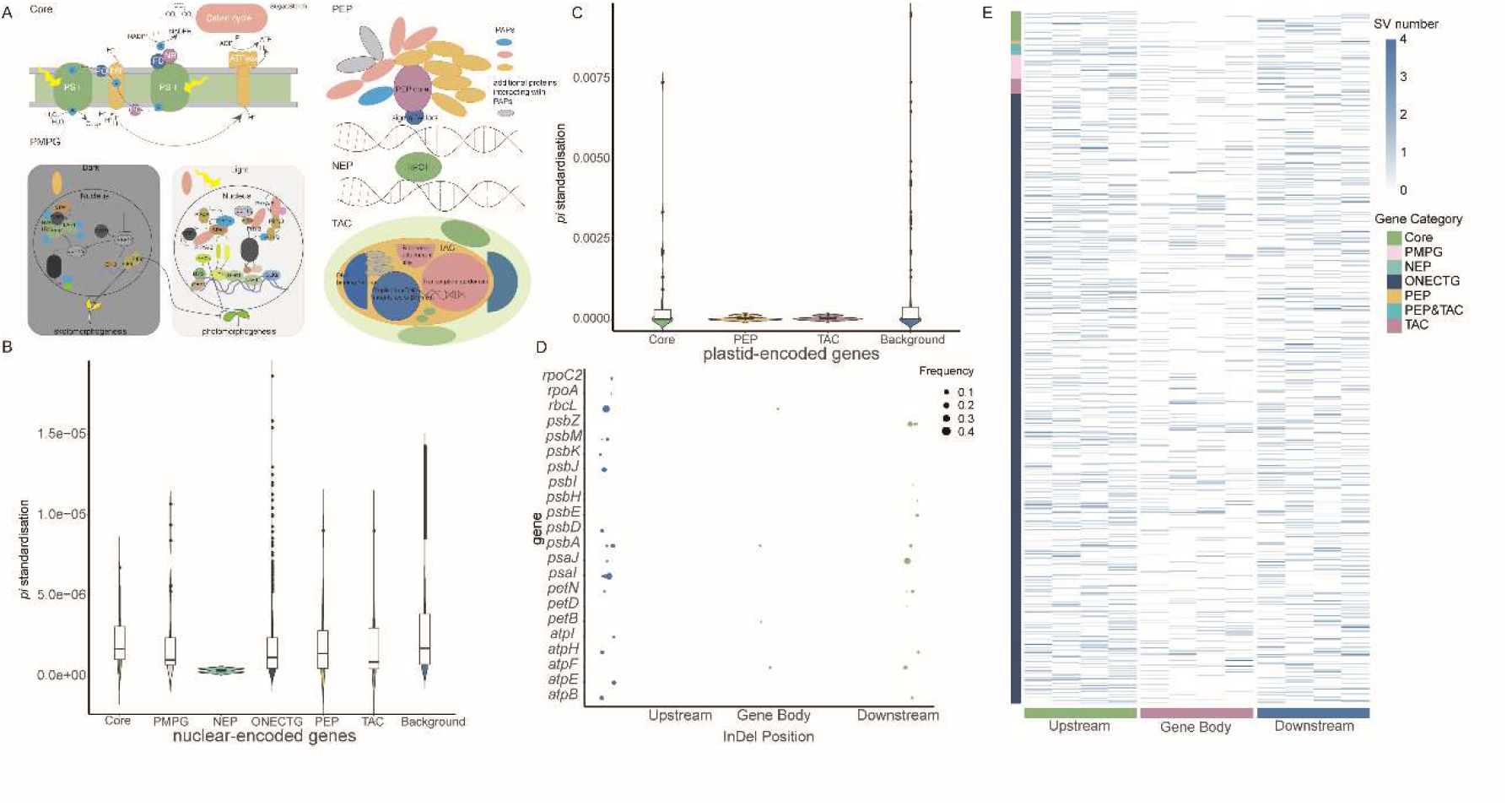
Rich genetic diversity in nuclear and plastid-encoded genes (ANPEG). (**A**) A schematic illustration of the photosynthesis associated pathways studied here. (**B**) The nucleotide diversity (*Pi*) of nuclear-encoded genes compared with the background. **(C)** The nucleotide diversity (*Pi*) of plastid-encoded genes compared with the background. (**D, E**) The number and frequency of Insertions/Deletions (InDels) and he number of structural variations (SVs) in ANPEG.

### Genetic regulation of the nuclear and plastid encoded genes (ANPEG)

To explore the genetic and transcriptomic diversity of ANPEG, we explored the *Arabidopsis thaliana* 1001 Genome dataset including sequenced genomes and transcriptomes from approximately 700 wild *Arabidopsis thaliana* accessions. Overall, the expression of ANPEG varied quantitatively among wild *Arabidopsis thaliana* accessions and showed intermediate kinship heritability (media 0.214, Fig. 3A and S3, Table S11). These results suggest that there is natural genetic variation and heritable expression variation for ANPEG in *Arabidopsis thaliana* wild population. Therefore, we applied expression quantitative trait locus (eQTL) mapping to dissect alleles underlying regulation of ANPEG. In total, we identified 332 eQTL associated with the expression of 376 genes (Fig 3B, Table S12). The majority of these eQTL were cis-eQTL (within 1 Mb of genes). However, one trans-eQTL hub located on chromosome 5: 22597100-22604800 bp was associated with the expression of 149 genes (Fig. 3B and 3D, Table S13). LD decayed rapidly at this region, and nearly all the association signals pointed to one candidate gene *AT5G55840* annotated in mRNA binding. The GG allele at this locus simultaneously increased the expression of 85.2% (127/149) ANPEG (Fig S4). Among these, several core genes, including *FD3, PSBO-2, PSBP-2*, and *PSBR* (Fig 3C) were found, suggesting that this region may regulate the expression of core genes and have an impact on photosynthetic efficiency.

**Fig 3.**
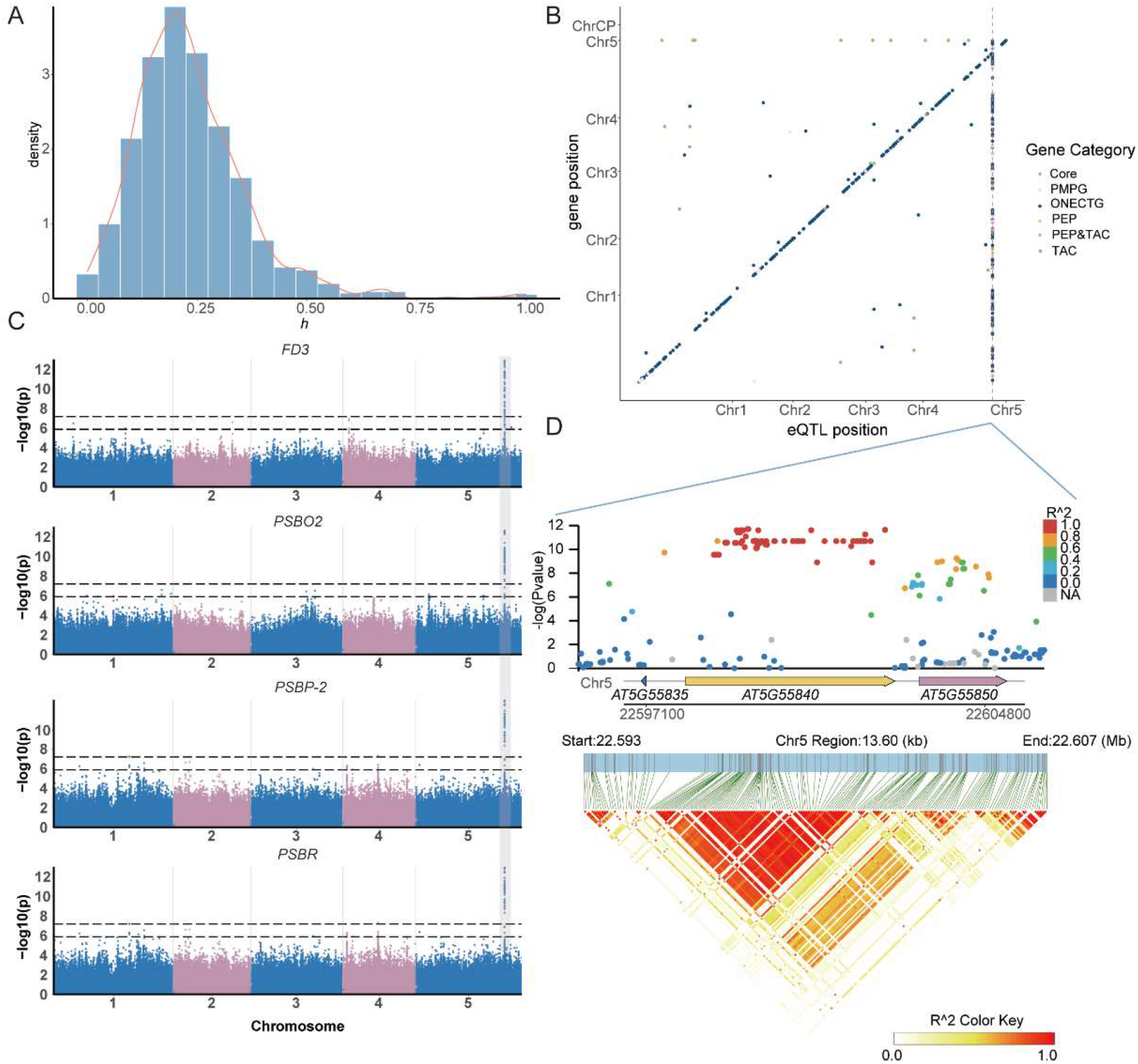
Genetic regulation of the nuclear and plastid encoded photosynthesis-associated genes. (**A**) Histogram of the kinship heritability (*h*^*2*^) for the expression of ANPEG. (**B**) A genome-wide map of eQTL. The x-axis shows the position of the eQTLs, the y-axis shows the location of the associated genes. (**C**) Manhattan plots of core genes associated with the trans-eQTL hub at chromosome 5:22597100-22604800 bp. (**D**) The LD (Linkage Disequilibrium) block plot of the trans-eQTL hub.

In summary, these results demonstrated abundant regulatory polymorphisms in wild *Arabidopsis thaliana* population, and identified a candidate gene that influences the expression of 39.6% (149/376) of the ANPEG.

### Transcriptome-wide association highlighted an important role of ANPEG expression variation in variation of Arabidopsis thaliana complex traits

To estimate the contribution of ANPEG expression variation to complex traits variation, we calculated the proportion of phenotypic variance (PVE) explained by the 1103 ANPEG using linear mixed model (Fig 4A, Table S14 and S15). Overall, these gene sets (less than 4.5%, median = 0.3%)) disproportionately explained a larger fraction of the phenotypic variance (0.8% to 23.5%, median= 8.7%). On average, background genes accounted for the largest proportion of phenotypic variance (32.1%), followed by ANPEG (23.5%), ONECTG (22.7%) and Core Genes (10.3%).

**Fig 4.**
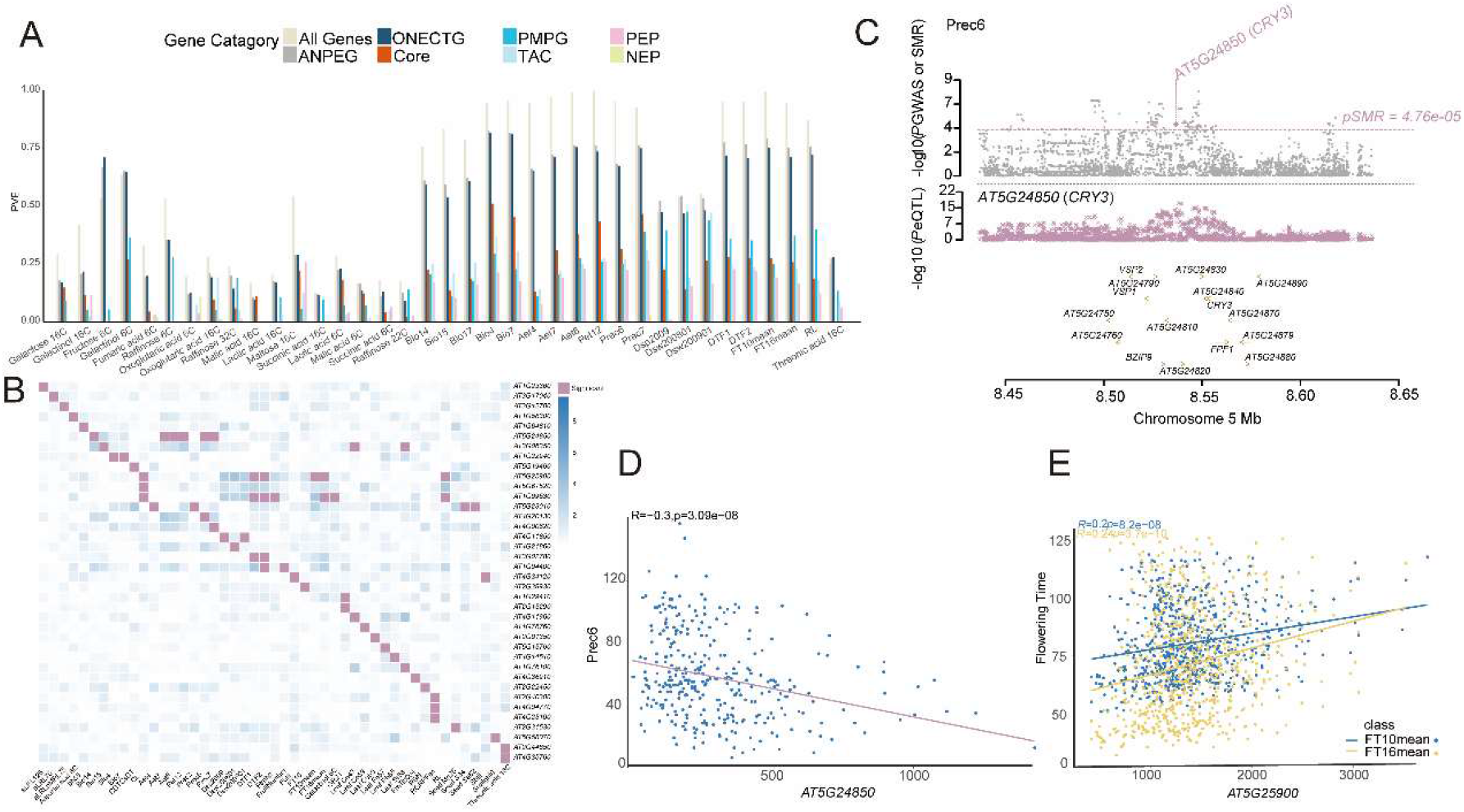
*Transcriptome-wide association* highlighted the important role of ANPEG expression variation in variation of *Arabidopsis thaliana* complex traits. (A) The proportion of phenotypic variance explained (PVE) by the expression variation of different gene categories. (B) Gene expression associated with 47 traits. The pink square represents the significant association. The blue square represents the gene’s *P-value* obtained from TWA analysis. (C) Prioritizing genes at a GWAS locus using SMR analysis. The top panel is the manhattan plot of Prec6 (Precipitation of June, mm) GWAS result. The middle panel is manhattan plot of *AT5G24850 (CRY3)* eQTL scan. The bottom panel is the gene annotation. (D) and (E) Expression-to-trait correlation maps for *AT5G24850* with Prec6 and *AT5G25900* with flowering time at 10 °C and 16°C.

To explore the role of ANPEG expression in complex trait variation, we performed a transcriptome-wide association analysis (TWA) analysis associating ANPEG expression with 475 traits downloaded from the AraPheno database and relevant publications (Table S14). 47 traits, including flowering time, ecological environmental factors, root morphology traits, mineral and ion content-related traits, and metabolite content traits exhibited significant associations with ANPEG expression (Table S16, Fig 4B, 4D,4E).

Furthermore, several genes with eQTL detected from previous section were associated with 27 traits. We, therefore, performed mendelian randomization analysis for casual inference. One gene (*AT5G24850, CRYPTOCHROME 3, CRY3*) with eQTL at chromosome 5:8538064 bp was associated with three ecological environmental traits (Prec6, Prec7, Aet8) (Fig. 4C, Table S17). This means that there was a single genetic variant regulating the expression of *AT5G24850 (CRY3)* and thereby affecting these three ecological environmental traits variation.

### eQTL associated with the expression of photosynthesis-core genes, and their association with the Arabidopsis thaliana complex traits

Ribulose-1,5-bisphosphate carboxylase/oxygenase (Rubisco) catalyzes a rate-limiting step in the Calvin cycle, transforming atmospheric carbon into a biologically useful carbon source. In all high plants, Rubisco is formed by large subunits (rbcL) and small subunits (rbcS). In *Arabidopsis thaliana*, small subunits are encoded by a gene family in the nucleus genome (*RBCS1A, RBCS1B, RBCS2B, RBCS3B*). Activation of Rubisco requires the removal of inhibitory sugar phosphates by Rubisco Activase (RCA). In our study, we detected three eQTL associated with the expression of *RBCS1B, RBCS2B*, and *RCA* (Fig. 5A, 5B and 5C). The detection of eQTL for these key Rubisco subunits highlighted the genetic regulation of photosynthetic efficiency through regulatory variants. Additionally, the eQTL of *RCA* was scientifically associated with fruit number and elevation (Fig 5D, 5E and S5). This suggests that variation of *RCA* may be associated with fruit Number (the total number of fruits per plant at the end of reproduction).

**Fig 5.**
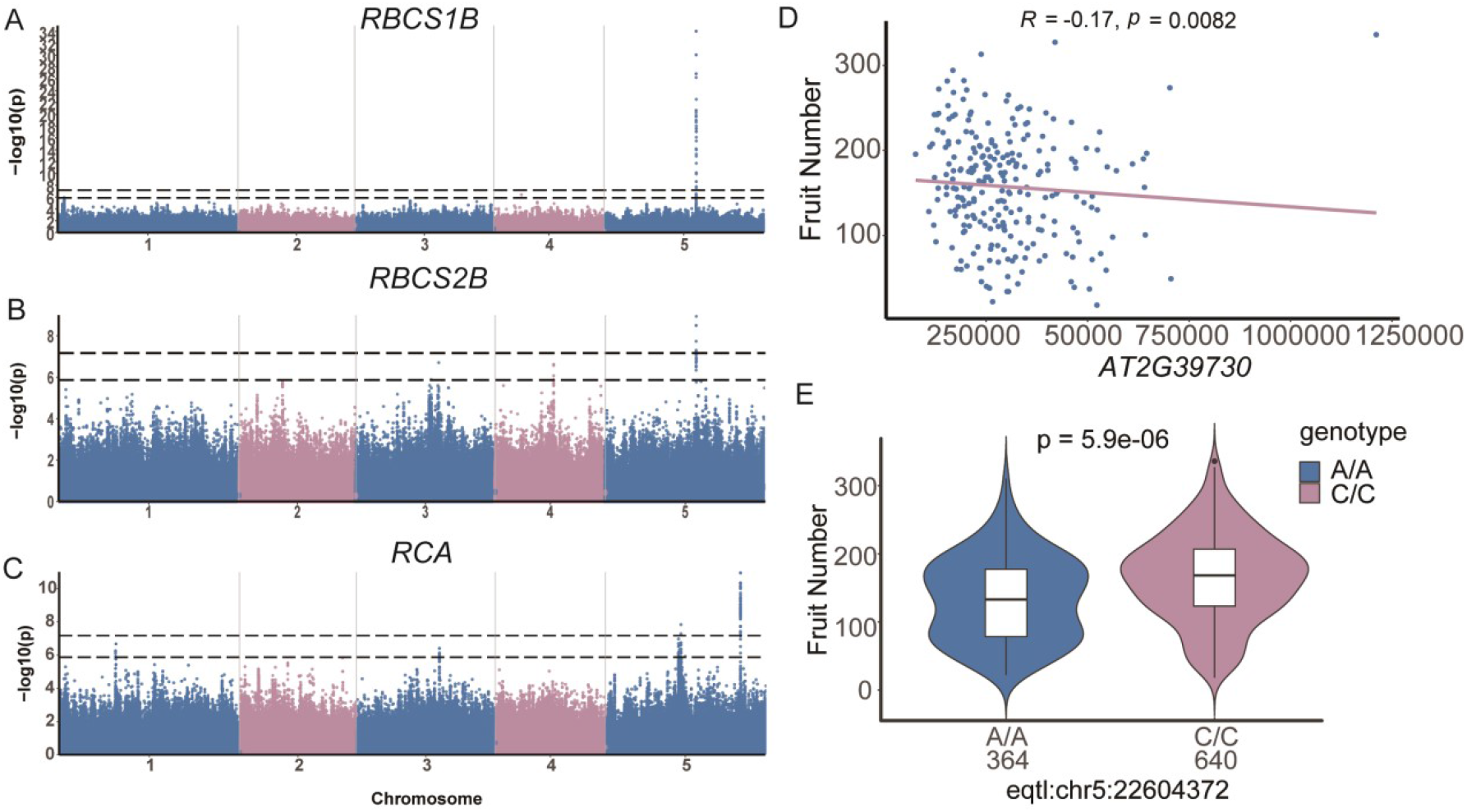
eQTL associated with the expression of core genes, and their association with the *Arabidopsis thaliana* complex traits. (**A**) (**B**) and (**C**) Manhattan plots of *RBCS1B, RBCS2B*, and *RCA*. (**D**) The association of the expression of *RCA* and fruit number. The coefficient *R* is derived from the Spearman’s rank correlation test. (**E**) The genotype to phenotype map for fruit number at Chromosome 5: 22604372 bp. *P-value* was obtained from the Wilcoxon test.

## Discussion

Previous studies revealed a limited diversity in core photosynthetic components and argued that genetic variation in photosynthesis likely occurs beyond the core photosynthetic process^32-34^. Here, by leveraging 28 *de novo* assembled chloroplast genomes, hundreds of accessions with whole genome and transcriptome data, we revealed abundant genetic diversity in plastid-encoded photosynthesis-associated genes (*psaJ, rbcL*, Fig S1, Fig 1D, Table S3). SVs and InDels were present in high frequently adjacent to key photosynthesis genes, such as *psaJ rbcL, ATPC1, PSBO-2, RBCS1B*, and etc. (Fig 2). These findings indicated there were extensive natural genetic variation in photosynthesis-core genes among worldwide *Arabidopsis thaliana* accessions, representing a significant reservoir to improve photosynthetic efficiency.

Here, we detected regulatory variation for 34.0% of the ANPEG through eQTL mapping. A trans-eQTL hotspot regulating the expression of 149 ANPEG (Fig 3) were found. Some of the eGenes (Genes with detectable eQTL) were in the core photosynthetic process. For example, RuBisCO acts as the rate-limiting enzyme in photosynthesis, thereby determining the overall efficiency of photosynthesis ^35^. We found three eQTL regulating the expression of *RBCS1B, RBCS2B*, and *RCA* (Fig 5). One of the eQTL was associated with increased fruit number and colocalized with a QTL associated with high elevation (Fig S5). Given that genetic variations of RuBisCO have been reported^8,36,37^, detection of regulatory variation in our study warrant further investigation on their role in local adaptation and their potential in boosting photosynthetic efficiency.

Here, we found that expression of ANPEG contributed to a large proportion of the phenotypic variation for a number of complex traits. Those traits ranging from metabolites, such as leaf sugar content, leaf ionomics traits, to complex fitness and reproductive traits, including flowering time, fruit number and environment factors at native range (Fig 4B, Table S14 and S15). Through mendelian randomization analysis, a single genetic variant was found to regulate the expression of *AT5G24850 (CRY3)* thereby impacting three precipitation and evapotranspiration-related traits (Fig 4C, Fig 4D, Table S17). These results highlighted the important contribution from natural genetic and transcriptomic variation in worldwide *Arabidopsis thaliana* photosynthesis associated pathways to complex adaptive traits variation.

Overall, our study revealed extensive natural genetic and transcriptomic variation of ANPEG among worldwide *Arabidopsis thaliana* accessions and demonstrated their role in complex traits variation and adaptation. Results from our study provides deeper insights into natural variation in photosynthesis associated pathways and their function in complex trait variation, paving new routes in characterizing and utilizing such variation in plant breeding.

## Materials and Methods

### Assembly of chloroplast genomes and nucleotide diversity calculation

Clean Pacbio Sequel II reads from 28 genomes were downloaded from Kang et al^38^. Chloroplast genome reads were extracted using Blast ^39^ and subsequently assembled using Canu v2.2 ^40^ to obtain a circular chloroplast genome sequence. The assembled sequence were validated by aligning to Col-0 (NC_000932.1) reference genome using minimap2 v2.26^41^ and then manually curated. A second round of validation was performed by mapping all the clean reads used for assembly back to our assembly using minimap2. Read depth was calculated from each assembly using SAMtools v1.18 ^42^ and visualized using ggplot2 in R (Fig S6). Nucleotide diversity was calculated with DnaSP v6 software^43^ and a scatter plot was made by the *circlize* v0.4.15^44^ package in R.

### List of ANPEG, RNA seq and phenotype dataset

We obtain the photosynthesis-core genes from the photosynthesis pathway annotation in the KEGG database (https://www.genome.jp/kegg/). The R package *UniportR* v2.3.0^45^ was utilized to obtain the other nuclear-encoded chloroplast-targeted genes. Other gene categories were sourced from related publications^46,47^. RNA-seq data of nuclear and chloroplast genomes were obtained for 726 accessions of *Arabidopsis thaliana* from NCBI (https://www.ncbi.nlm.nih.gov/geo/, GEO accession ID: GSE80744). The downloaded read counts for nuclear genome data were already normalized. The raw counts for chloroplast genome data were normalized to TPM (transcripts per million) using *GenomicFeatures* v3.18^48^ package in R. 475 traits of *Arabidopsis thaliana* were downloaded from the AraPheno database (https://arapheno.1001genomes.org/phenotypes/) and some reasearches^49,50^.

### Variants Calling and summary statistics

The genotype data in the nuclear genomes was downloaded from the *Arabidopsis thaliana* 1001 Genome Project website (https://1001genomes.org/data/GMI-MPI/releases/v3.1). SV (Structure Variant) in the nuclear genomes was downloaded from Kang et al^38^, SVs were then filtered with parameters setting for minor allele frequency more than 0.03. Fastq-dump in the SRA toolkit (https://ftp-trace.ncbi.nlm.nih.gov/sra/sdk/2.8.0/) was used to convert the SRA data to fastq files. Raw reads were first quality controlled using fastp v0.23.4 ^51^ with default parameters, then SAMtools v1.18^42^ and BCFtools v1.9^52^ were used for variant calling for the chloroplast genomes. SNPs and InDels were filtered using VCFtools v0.1.16^53^ for keeping bi-allelic, deletion rates below 80%, and minor allele frequency more than 0.03.

Nucleotide diversity for the nuclear genomes was calculated using VCFtools, and nucleotide diversity in the chloroplast genomes was calculated using *Popgenome v*2.7.5 package^54^ in R. To compare Nucleotide diversity (*Pi*) between ANPEG and the genetic diversity of all genes of the reference genome (Col-0)(background), *Pi* was initially adjusted for gene length, then a Wilcoxon test was performed in R. Here, we using the first 90% *Pi* of all genes to represent the distribution trend of genetic diversity in all genes to avoid the outliers disturbing the Wilcoxon test. The frequency and number of SVs and InDels were counted using VCFtools. SVs located 5k bp upstream and downstream of nuclear-encoded genes were summarized, and InDels located 500 bp upstream and downstream of plastid-encoded genes were summarized.

### Expression quantitative trait locus (eQTL) mapping

eQTL mapping was performed using the R package GenABEL v1.8.0 ^55^ using the *mmscore* function. The genome-wide significant threshold (Suggestive *P-value*:1.35E-6, Significant *P-value*:6.74E-8) of eQTL mapping was estimated using gec.jar v0.2 ^56^ in Java, which adjusts for the effective number of SNPs based on linkage disequilibrium. The Linkage Disequilibrium (LD) block of the trans-eQTL hub was estimated and visualized using LDBlockShow v1.40^57^.

### Estimating the contribution from transcriptome to Arabidopsis thaliana complex traits variation and transcriptome-wide association analysis

A mixed linear model was used to estimate the proportion of phenotypic variance explained by ANPEG transcripts.

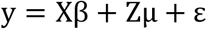

where y is an n × 1vector of phenotype values with n being the sample size, and Z is an n × m matrix of standardized gene expression measures of all m genes. X is an n × 1 vector of 1. In this model, β are fixed effects, μ and ε are random effects with 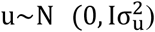 and 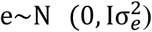.

Transcriptome-wide association analysis (TWAS) was conducted by mixed linear model approaches (MOA) using OSCA 0.46.1^58^. The Benjamin and Hochberg FDR threshold (FDR < 0.05) was used as the gene-wide significance threshold.

### Summary data-based Mendelian randomization analysis

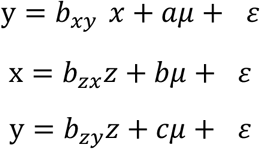

Let y be a phenotype,x be gene expression,z be a genetic variant^59^,*b*_*xy*_ be the effect size of x on y, *b*_*zy*_ be the effect of z on y. *b*_*xy*_ can be estimated and tested using summary-level data which is called the summary data–based Mendelian randomization (SMR) method, and we distinguished pleiotropy from linkage using the HEIDI (heterogeneity in dependent instruments) test in SMR v1.3.1 to prove there is a single variant affecting both phenotype and gene expression (*P*_*HEIDI*_ > 0.05).

